# BtToxin_Digger: a comprehensive and high-throughput pipeline for mining toxin protein genes from *Bacillus thuringiensis*

**DOI:** 10.1101/2020.05.26.114520

**Authors:** Hualin Liu, Jinshui Zheng, Dexin Bo, Yun Yu, Weixing Ye, Donghai Peng, Ming Sun

## Abstract

*Bacillus thuringiensis* (Bt) which is a spore-forming gram-positive bacterium, has been used as the most successful microbial pesticide for decades. Its toxin genes (*cry*) have been successfully used for the development of GM crops against pests. We have previously developed a web-based insecticidal gene mining tool BtToxin_scanner, which has been proved to be the most important method for mining *cry* genes from Bt genome sequences. To facilitate efficiently mining major toxin genes and novel virulence factors from large-scale Bt genomic data, we re-design this tool with a new workflow. Here we present BtToxin_Digger, a comprehensive, high-throughput, and easy-to-use Bt toxin mining tool. It runs fast and can get rich, accurate, and useful results for downstream analysis and experiment designs. Moreover, it can also be used to mine other targeting genes from large-scale genome and metagenome data with the addition of other query sequences.

**Availability and Implementation:** The BtToxin_Digger codes and instructions are freely available at https://github.com/BMBGenomics/BtToxin_Digger. A web server of BtToxin_Digger can be found at http://bcam.hzau.edu.cn/BtToxin_Digger.

## 1 Introduction

The toxins produced by *Bacillus thuringiensis* (Bt) have insecticidal activity against many agricultural and forestry pests, so they are widely used in the development of biopesticides and GM insect-resistant crops. Bt products represent more than 60% of the biopesticide market (Siegwart *et al*., 2015). Crystal protein (Cry) produced by Bt as the major toxin can kill insects from many orders including Lepidoptera, Diptera, and Coleoptera, etc. The *cry* gene is one of the most important genes used for the development of genetically modified (GM) crops targeting insect pests. From 1996 to 2016, the planting of Bt maize and cotton had delivered $50.6 billion and $54 billion of extra farm income, respectively (G Brookes and Barfoot, 2018). Due to the importance of Bt toxins, many researchers and companies have been working on the discovery of new toxin genes (Sanahuja *et al*., 2011). Other toxins with insecticidal activity produced by Bt include Cyt (Cytotoxic toxin protein) and Vip (Vegetative insecticidal protein), etc (Palma *et al*., 2014). Previously, we developed an on-line tool BtToxin_scanner to predict Crys encoding genes from Bt genome sequences (Ye *et al*., 2012). It can handle several assembled genomes every time and provide useful comparative results between the precited toxin and with known ones. During the past 7 years, it was widely used by researchers worldwide (Méric *et al*., 2018; Prado *et al*., 2014; Ruan *et al*., 2015; Zheng *et al*., 2017). Here we re-designed the previous tool to provide a novel, high-throughput, and local software BtToxin_Digger which can be directly used to handle large-scale genomic and metagenomic data to predict all kinds of putative toxin genes. It also generates comprehensive and readable results to facilitate the downstream sequence analysis or experiment design (Figure 1).

**Figure 1.**
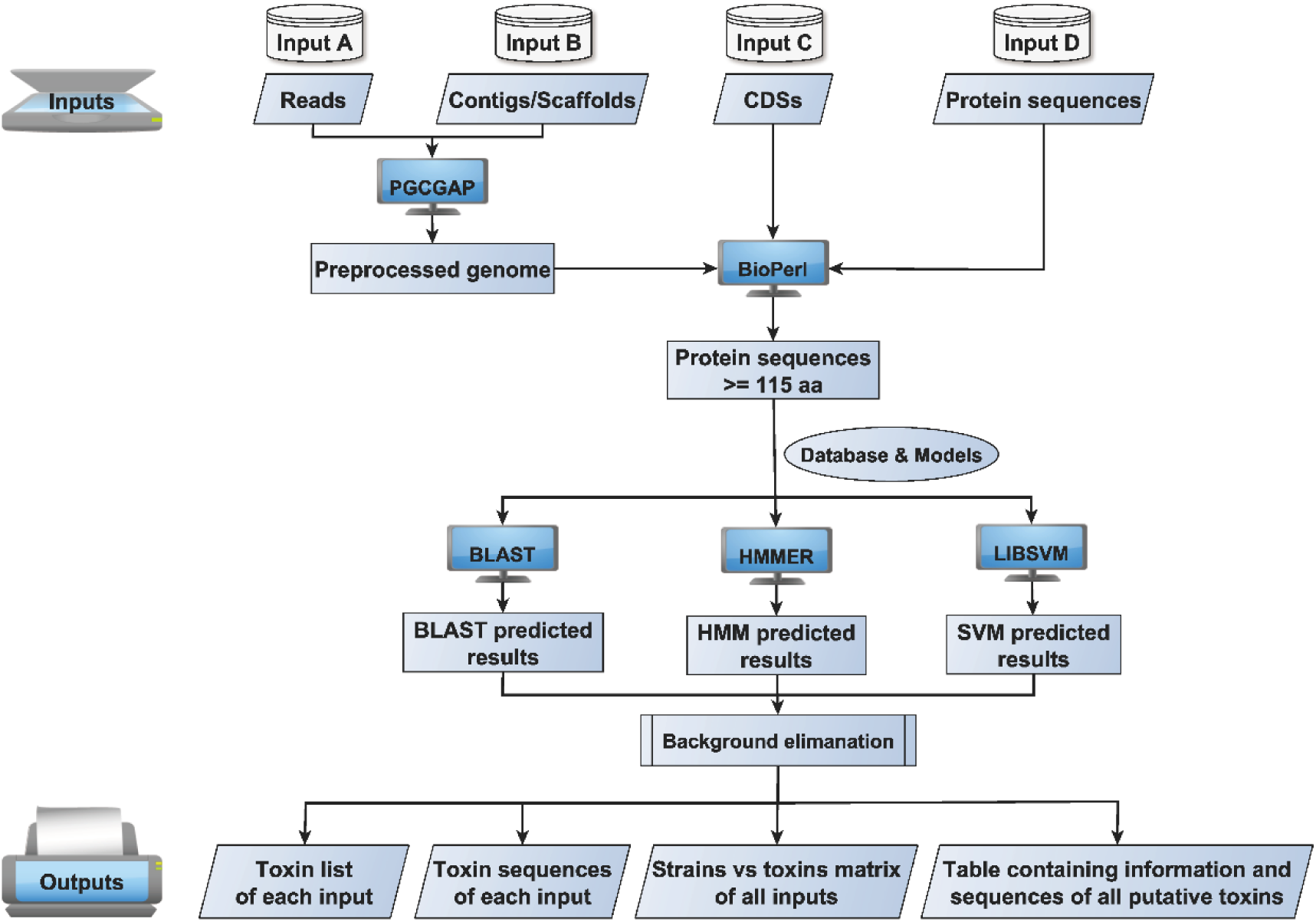
A diagram of the BtToxin_Digger pipeline.

## 2 Methods

The tool accepts multiple forms of input data including Reads (pair-end reads, long-reads, or hybrid-reads), genomic or metagenomic assemblies, coding sequences (CDSs), and protein sequences. PGCGAP (Liu *et al*., 2020) was used for genome assembly and pretreatment. ORFs finding and translation are performed by BioPerl (Stajich *et al*., 2002). All protein sequences with a length above 115-aa are searched against the database and trained models by BLAST (Camacho *et al*., 2009), HMMER (Eddy, 2011), and LIBSVM (Chang and Lin, 2011), respectively. After that, the candidate proteins are blasted against a background database to filter out the false-positive records. Then several Perl scripts are used to parse the results to get the putative target protein genes.

## 3 Results

BtToxin_Digger can be easily installed on Linux, macOS, and Windows Subsystem for Linux (WSL) platforms by the conda package manager (Grüning *et al*., 2018) or docker container. We tested BtToxin_Digger on a laptop with an Intel CPU containing 8 threads of GHz-2.50 and 16 GB memory. It took 14 minutes to process the 1.3-Gbp raw reads to get the results. Moreover, it just takes less than one minute to finish the whole analysis when the other three inputs were provided. BtToxin_Digger can also be used to mine other interesting protein genes with the replacement of the Bt toxin database by other target sequences. We also developed a webserver http://bcam.hzau.edu.cn/BtToxin_Digger for users with less data (assembled genomes, amino acid sequences and coding sequences only) to analize.

We compared BtToxin_Digger with the existing tool BtToxin_scanner (Ye *et al*., 2012) and CryProcessor (Shikov *et al*., 2020). As can be seen from Table 1, BtToxin_ Digger adopts more mining methods, supports more types of input files and toxins, and gets more friendly output results. Compared with the other two software, it is more suitable for large-scale toxin gene mining, and at the same time, it can easily implement the high-throughput analysis.

**Table 1.** Comparation of BtToxin_Digger, BtToxin_scanner and CryProcessor.

| Tool | Main methods used | Supported inputs | Supported toxins | Outputs | Flux |
| --- | --- | --- | --- | --- | --- |
| BtToxin_Digger | Blast, HMM, SVM | Illumina/Pacbio/Oxford reads, assembled genomes, protein sequences, coding sequences, ORFs | Cry, Cyt, Vip, Other toxins | Toxin list file and sequences file for each input, a matrix file describes all strains vs. all toxins, an integrated file contains sequences and information of all inputs | Unlimited number of inputs with a one-line command |
| BtToxin_scanner | Blast, HMM, SVM | Assembled genomes, protein sequences, ORFs | Cry | Toxin list file and sequences file for each input | One submit at a time |
| CryProcessor | HMM | Illumina reads, representing genome assembly graph files, protein sequences | Three-domain Cry | A directory containing multiple files for each input | Unlimited number of inputs with additional shell scripts |

### Practice with the sample dataset

We also provide the sample dataset to demonstrate the usage of BtToxin_Digger (Supplementary File 1). To use this tool, users should install it on their computers and have a preliminary understanding of Linux. Users can refer to the protocol (Liu *et al*., 2020) to build their bioinformatics analysis platform and refer to https://github.com/BMBGenomics/BtToxin_Digger#installation to install BtToxin_Digger. We also prepared a webpage (https://github.com/liaochenlanruo/pgcgap/wiki/Learning-bioinformatics) for users without Linux skills to learn the basic Linux commands. Because the reads file is too large for upload and download, here we only demonstrate the running method of assembled genome, protein sequences, and coding sequences. Users can visit https://github.com/BMBGenomics/BtToxin_Digger#examples for more information.

Step 1. Download the Example dataset (Supplementary File 1) and unzip files.

Step 2. Open a terminal and enter the directory.

~~~
cd ExampleDataset
~~~

Step 3. Processing assembled genomes

~~~
BtToxin_Digger --SeqPath ./Genome --SequenceType nucl --Scaf_suffix .fas --threads 4
~~~

Step 4. Processing protein sequences

~~~
BtToxin_Digger --SeqPath ./AAs --SequenceType prot --prot_suffix .faa --threads 4
~~~

Step 5. Processing coding sequences

~~~
BtToxin_Digger --SeqPath ./CDSs --SequenceType orfs --orfs_suffix .ffn --threads 4
~~~

The running results are stored in Supplementary File 2. *.list: toxin list of each strain; *.gbk: toxin sequences in Genbank format of each strain; Bt_all_genes.table: a matrix describes Strains vs. Toxins; All_Toxins.txt: a table containing all information and sequences of all toxin genes. See Supplementary Table 1 for details.

## Supporting information

Example Dataset

Example Results

## References

Camacho, C., et al. (2009) BLAST+: architecture and applications. BMC Bioinformatics, 10, 421.

Chang, C.-C. and Lin, C.-J. (2011) LIBSVM: A library for support vector machines. ACM Trans. Intell. Syst. Technol., 2, Article 27.

Eddy, S.R. (2011) Accelerated Profile HMM Searches. PLoS Comp Biol, 7, e1002195.

G Brookes and Barfoot, P. GM crops: global socio-economic and environmental impacts 1996-2016. UK: PG Economics Ltd; 2018.

Grüning, B., et al. (2018) Bioconda: sustainable and comprehensive software distribution for the life sciences. Nat Methods, 15, 475–476.

Liu, H., et al. (2020) Build a bioinformatics analysis platform and apply it to routine analysis of microbial genomics and comparative genomics. Protocol Exchange, DOI: 10.21203/rs.2.21224/v2.

Méric, G., et al. (2018) Lineage-specific plasmid acquisition and the evolution of specialized pathogens in *Bacillus thuringiensis* and the *Bacillus cereus* group. Mol Ecol, 27, 1524–1540.

Palma, L., et al. (2014) *Bacillus thuringiensis* toxins: an overview of their biocidal activity. Toxins, 6, 3296–3325.

Prado, J.R., et al. (2014) Genetically Engineered Crops: From Idea to Product. Annu Rev Plant Biol, 65, 769–790.

Ruan, L., et al. (2015) Are nematodes a missing link in the confounded ecology of the entomopathogen *Bacillus thuringiensis*? Trends Microbiol, 23, 341–346.

Sanahuja, G., et al. (2011) *Bacillus thuringiensis*: a century of research, development and commercial applications. Plant Biotechnol J, 9, 283–300.

Shikov, A., et al. (2020) No More Tears: Mining Sequencing Data for Novel Bt Cry Toxins with CryProcessor. Toxins, 12, 204.

Siegwart, M., et al. (2015) Resistance to bio-insecticides or how to enhance their sustainability: a review. Front Plant Sci, 6, 381.

Stajich, J.E., et al. (2002) The Bioperl toolkit: Perl modules for the life sciences. Genome Res, 12, 1611–1618.

Ye, W., et al. (2012) Mining new crystal protein genes from Bacillus thuringiensis on the basis of mixed plasmid-enriched genome sequencing and a computational pipeline. Appl Environ Microbiol, 78, 4795–4801.

Zheng, J., et al. (2017) Comparative Genomics of *Bacillus thuringiensis* Reveals a Path to Specialized Exploitation of Multiple Invertebrate Hosts. MBio, 8, e00822–00817.

