## Supplementary Table 1 for "BtToxin_Digger: a comprehensive and high-throughput pipeline for mining toxin protein genes from *Bacillus thuringiensis*"

Supplementary Table 1. Description of “All_Toxins.txt”

| Header | Description |
|---|---|
| Strain | The name of your input |
| Protein_id | The protein ID |
| Protein_len | The length of protein sequence |
| Strand | Positive or negative strand where the gene comes from |
| Gene location on scaffold | Gene coordinates on the genome |
| SVM | Is the protein predicted by SVM |
| BLAST | Is the protein predicted by BLAST |
| HMM | Is the protein predicted by HMM |
| Hit_id | The subject sequence ID |
| Hit_length | The length of subject sequence |
| Aln_length | alignment length (sequence overlap) |
| Query start-end | Start and end of alignment in query |
| Hit start-end | Start and end of alignment in subject |
| Identity | Percentage of identical matches |
| Evalue of blast | Expect value of BLAST |
| Hmm hit | The subject model ID |
| Hmm hit length | The length of subject model sequence |
| Evalue of Hmm | Expect value of HMM |
| Nomenclature | Bt nomenclature containing 4 Ranks |
| Endotoxin_N | Whether the Cry protein contain Endotoxin_N domain |
| Endotoxin_M | Whether the Cry protein contain Endotoxin_M domain |
| Endotoxin_C | Whether the Cry protein contain Endotoxin_C domain |
| Endotoxin_mid | Whether the Cry protein contain Endotoxin_mid domain |
| Toxin_10 | Whether the Cry protein contain Toxin_10 domain |
| ETX_MTX2 | Whether the Cry protein contain ETX_MTX2 domain |
| Gene sequence | The nucleotide sequence of the toxin |

| Header | Description |
| --- | --- |
| Protein sequence | Amino acid sequence of the toxin |
| Scaffold sequence | The scaffold sequence where the toxin gene is located |
